# A multi-stage, delay differential model of snow crab population dynamics in the Scotian Shelf of Atlantic Canada

**DOI:** 10.1101/2023.02.13.528296

**Authors:** Jae S. Choi

**Affiliations:** Bedford Institute of Oceanography, Fisheries and Oceans Canada, Challenger Dr., Dartmouth, Nova Scotia, B2Y 4A2, Canada

## Abstract

Snow crab (*Chionoecetes opilio*) are cold-water stenotherms in the northern hemisphere. As they are long-lived and have a complex life history, developing an operational model of population dynamics has been a challenge, especially in the context of an ever increasing and varied human footprint upon nature. Here we review past efforts at understanding the population dynamics of snow crab in an environmentally and spatiotemporally heterogeneous area, the Scotian Shelf of the northwest Atlantic of Canada. We address these difficulties with a moderately complex multi-stage, delay differential model and parameterize it leveraging Bayesian techniques. Operational *solutions* were stable and reasonable and permitted inference upon the intra-annual dynamics of snow crab. Further, a concept of a *Fisheries footprint*, akin to instantaneous fishing mortality rate, can be elucidated that directly addresses the conceptual impact of a fishery upon a non-stationary population. The approach is promising. The model suggests additional processes need to be accounted. We hypothesize that seasonal, interannual movement and spatiotemporally structured predation are key processes that require further attention. However, as computational costs are significant, these additional processes will need to be parameterized carefully.

## 1 Introduction

Snow crab (*Chionoecetes opilio*) are large Crustaceans, exploited primarily as a food source. They are also used as fertilizer, bait in other fisheries and for the glucosamine polysaccharide derived from chitin, known as chitosan. Chitosan is widely used in medicine as an agent to slow bleeding from wounds [1, 2], agriculturally as natural fungicides and bactericides [3], plastic replacement [4] and even as a battery electrolyte [5]. In North America, including in the area of this study, the Scotian Shelf of the northwest Atlantic of Canada (Figure 1), the largest of males (> 95 mm carapace width) are preferentially captured due to their higher meat-yield and consumer preferences of larger claws, a trait that occurs only at the terminal molt to maturity. This market-driven protection for the female reproductive crab which never reaches such sizes and the smaller immature crab represents a form of built-in protection for the population which spans 10 years of age or more, from larval release; it helps to offset otherwise heavy exploitation pressures of mature males by fishers with advanced technological and historical knowledge of their environment and the species’ aggregation patterns. Currently, every known population of snow crab is exploited by humans. As such, it is imperative that exploitation occurs in a responsible and sustainable manner.

**Figure 1:**
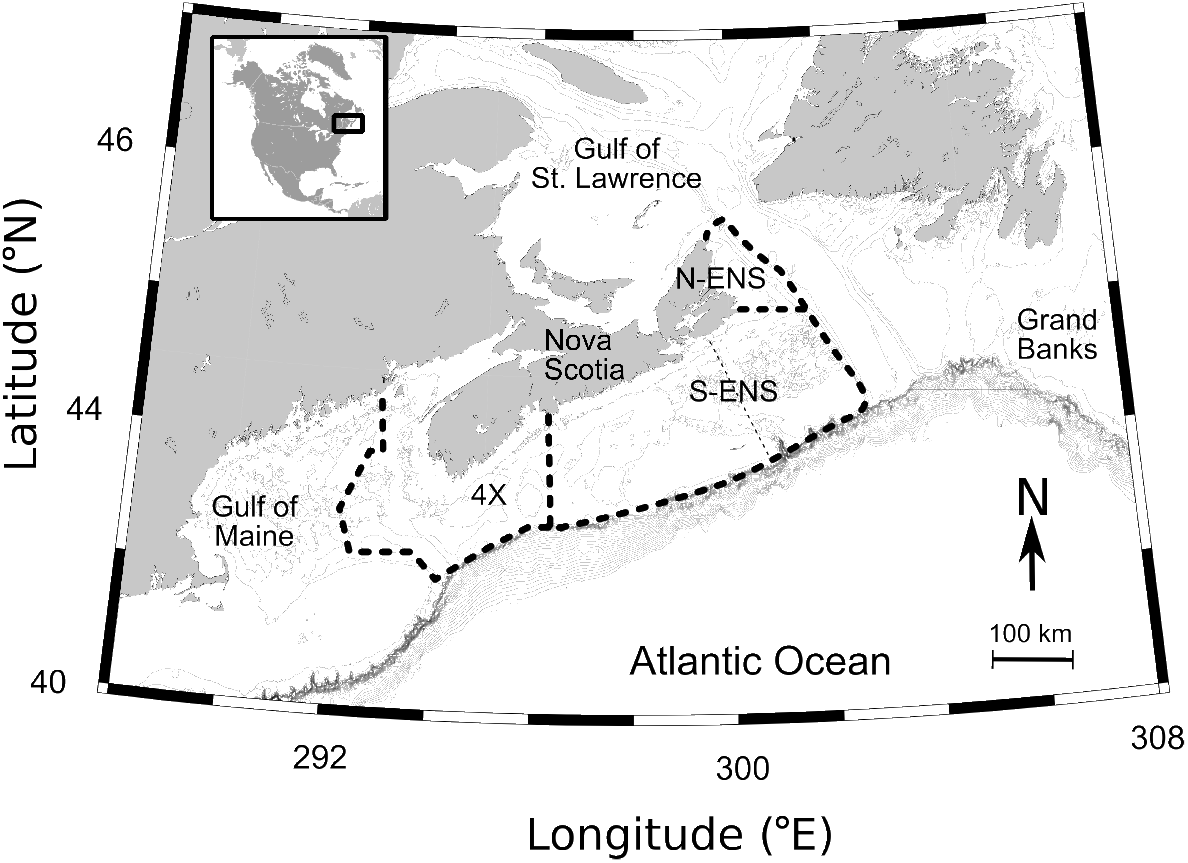
The area of interest in the Scotian Shelf of the northwest Atlantic Ocean, Canada. This area is at the confluence of the Gulf Stream from the south and south east along the shelf edge, Labrador Current and St. Lawrence outflow from the north and north east, as well as a nearshore Nova Scotia current, running from the northeast. It is hydro-dynamically complex due to mixing of cold, low salinity water from the north with the warm saline water from the south. Shown also are the managed Crab Fishing Areas divided by thick dashed lines: NENS (North-Eastern Nova Scotia), SENS (South-Eastern Nova Scotia), CFA 4X (Crab Fishing Area 4X).

Unfortunately, characterizing snow crab population dynamics is remarkably difficult (Figure 2) because they are long lived, have a complex life history, and large vertical and horizontal shifts in spatial distributions as they grow older (ontogenetic shift in habitat). Functionally, they participate in the pelagic and subsequently benthic ecosystems and associated nutrient and carbon cycles. Some of the notable life history features of snow crab that make them so interesting but difficult to model, include: sexual dimorphism with mature males being much larger than mature females; pelagic larval stages and benthic pre-adolescent and adult stages; semi-annual and annual molts depending upon size, age and environmental conditions; skipping moults if conditions are poor; terminal molt to maturity; and longevity up to 15 years. They also have a narrow range of temperature preferences [6, 7, 8]. They are thought to avoid temperatures above 7^*°*^*C*, as metabolic costs have been shown to exceed metabolic gains above this temperature in laboratory studies [6]. Smaller crab and females also have differences in thermal preferenda [9]. Further, snow crab are generally observed on soft mud bottoms; with smaller-sized and molting crabs showing preference for more complex (boulder, cobble) substrates, presumably as they afford more shelter [10, 11]. This life history complexity induces complexity (non-random structure) in their dynamics and spatial distributions.

In the area of study, the continental shelf of the northwest Atlantic Ocean (Figure 2), snow crab are further exposed to high bottom temperature variability due to the confluence of a number of oceanic currents: the warm Gulf Stream, the cold Labrador current, low salinity outflow from the St. Lawrence river and the cold coastal Nova Scotia current. In this region, snow crab are generally observed between depths of 50 to 300 m and between temperatures of −1 to 11^*°*^*C* [12]. As the focal area is thermally complex, their spatial distributions can fluctuate seasonally and annually. The additional factors of global rapid climate and ecosystem change confounds make this understanding even futher.

**Figure 2:**
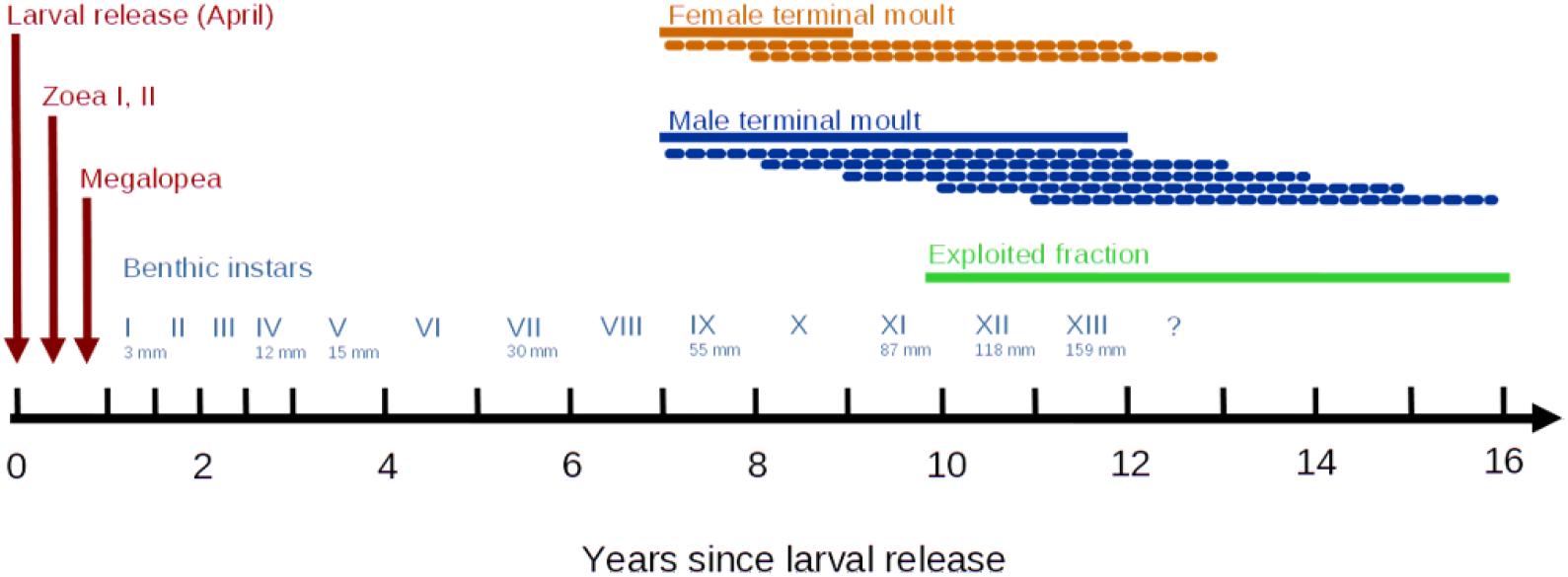
Life history patterns of snow crab and approximate timing of the main life history stages since larval release of snow crab and size (measured as carapace width; mm) and instar (Roman numerals). Size and timings are specific to the area of study and likely vary with regional environmental conditions, food availability and genetic variability. Brooding time is variable and is between 1 and two years. Initiation of terminal molt to maturity (orange and blue lines) also vary in timing. After initiation, longevity is thought to be about 5 to 6 years (dashed lines). Green solid line identifies approximate size and age of males exploited by humans.

**Figure 3:**
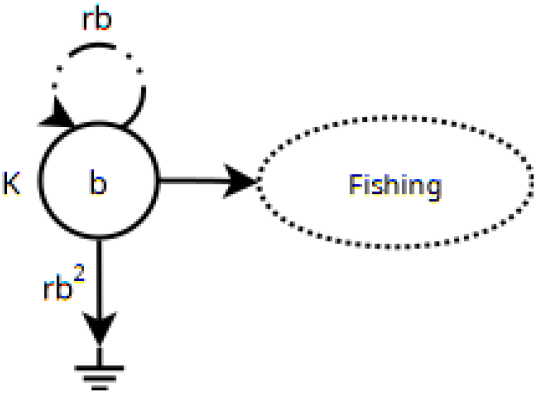
A graphical representation of Model 1 (simple logistic; eq. 1). Here, b = BK^−1^ is a non-dimensional number that ranges from (0, 1) that represents the biomass B after being scaled to K. The loop rb identifies the growth rate and the loss term rb^2^ identifies the quadratic increase in “mortality” as b → 1. Fishing is also scaled to K, such that: 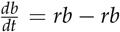. b – f = birth – death – fishing. As such, rb2 represents the non-fishing related mortality that is density dependent.

Snow crab eggs are brooded by their mothers for up to two years, depending upon ambient temperatures as well as food availability (which also depends upon temperature as it influences overall system primary and secondary production) and the health condition and maturity status of the mother (up to 27 months in primiparous females (first breeding event) event; and up to 24 months in multiparous females (second and subsequent breeding events) [13]. More rapid development of eggs, from 12 to 18 months, has been observed in other areas [14, 15]. Over 80% of the female snow crab on the Scotian Shelf are estimated to follow an annual cycle, possibly due to extended periods of exposure to warmer temperatures [8]. A primiparous female of approximately 58 mm carapace width produces between 35,000 to 46,000 eggs, which are extruded between February and April [13]. Multiparous females are thought to be more fecund, with more than 100,000 eggs being produced by each female. Eggs are hatched from April to June when larvae are released. This pelagic larval stage lasts for three to five months (Zoea stages I and II) during which snow crab feed upon zooplankton. Pelagic stages seem to have highest abundance in October and so may begin settling in January [8]. Thereafter, they settle closer to the ocean bottom in their Megalopea stage. Very little is known of survival rates at these early life stages, but they are thought to depend highly upon temperature [7].

There have only been a small number of attempts at modeling the dynamics of snow crab populations. Being long-lived, sexually dimorphic, pelagic and benthic organisms, usually living far offshore, they are not easily surveyed. Most population assessments approach abundance estimation with a fishery-or survey-based relative index of a narrow segment of the population (the fishable component). Further, aging of snow crab is not possible due to a lack of retention of calcified body parts through molts. This has encouraged adoption of size-based models [16, 17, 18], often with strong and questionable assumptions. For example, [16] assume the male fishable component is the spawning stock biomass, though of course it is the females that are the reproductive component and completely unexploited with a shorter life span. Similarly, [19] implicitly assumes that the fishable biomass regenerates itself by application of a biomass dynamics model. Importantly, sex and size-selectivity of sampling gear are almost always assumed to be a constant or some smooth monotonic function of body size, and so static across time and space or known without bias. These are significant and biologically problematic assumptions. The problems encountered are fundamental issues shared with all other attempts at assessment. Specifically, they are the interplay between snow crab life history and behavior; our inability to observe them without bias due to sampling design being non-random with respects to the environmental factors controlling their distribution and abundance; and the sampling machinery (mesh size, trawls that cannot access rugose areas) that can only observe a small fraction of the population.

This paper identifies a novel approach that overcomes some of these difficulties in modelling snow crab, in particular, by incorporating observation error across sex and stage-structure (often called a *latent* or *state-space* model) and non-stationary habitat variability to simultaneously model population dynamics and infer population parameters. We examine its utility and demonstrate the mechanism by which we can make these and more complex models operationally viable through the use of efficient and modular computing that simultaneously models and infers parameters by leveraging the **Julia** programming environment [20] and the supporting **Turing** library [21] for Bayesian parameter inference and the **DifferentialEquations** library [22] for dynamical modeling. This examination is conducted in the Scotian Shelf Ecosystem (Figure 1) where the variability of ocean climate is known to be high [23, 24] and where we also have a consistently sampled population since the late 1990s. The principal conclusion of this study is that the novel proposed model is informative and demonstrates utility in understanding snow crab dynamics, integrating all available information and versatility in incorporating/aggregating ecosystem effects through its inlfuence upon habitat viability. It is sufficiently flexible in approach to permit other ecosystem factors such as classical inter-specific interactions and movement, which are planned for future studies.

## 2 Methods

Data collection and subsequent index estimation are described in [25, 26, 9]. In short, sampling in an unbiased manner under complex hydrodynamic conditions is particularly difficult as the random (spatially) stratified sampling that is usually adopted to account for such variability, fail to do so, due to the large dynamic (i.e.,spatiotemporal) structures operating on scales comparable to the domain. A model-based approach (*Conditional AutoRegressive Spatiotemporal Models*, **CARSTM**, https://github.com/jae0/carstm), a simple extension of Generalized Linear Models that accounts for spatiotemporal autocorrelated random effects in a Bayesian context and computed with INLA [27], are used to address the estimation problem. More specifically, biases induced by differential spatiotemporal variability in habitat for differing life stages which, upon aggregation across space, permits an informative index of abundance comparable across time (years). This study focuses upon the indices of abundance derived from this process for the period from 1999 to 2022. Importantly, no survey was conducted in 2020 due to Covid-19 related uncertainties. The index generation method being Bayesian, imputes these estimates, but variability attached to this year is large. These aggregate timeseries data and supporting **Julia** models used in this paper are available at: https://github.com/jae0/dynamical_model/tree/master/snowcrab.

The well understood *phenomenological* single component model of Verhulst [28, 29] is the usual starting point for most models and has been used to model snow crab for many years [19, 30]:

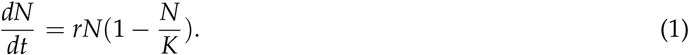

It is parameterized by an intrinsic rate of increase (*r*) representing the maximum exponential rate of increase and carrying capacity (*K*) the upper threshold. When numerical abundance *N* → 0, the loss term also 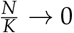, and so 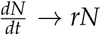. When *N* → *K*, then 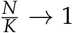 and so the loss rate approaches *rN* and the overall rate of change approaches zero 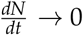. Parenthetically, this can be simplified further by dividing both sides by *K* to give: 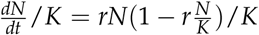. By focusing upon 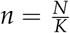 as a non-dimensional number ranging from (0,1), this becomes: 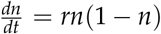. For parameter estimation/inference, this form is useful as it is simpler and the magnitudes of variables are mostly similar, with the exception of *K*. This renders beneficial properties to numerical optimization procedures. The sigmoid nature of the equation has rendered it a frequently encountered model that shows versatility in phenomenologically describing population dynamics. Its discrete form is particularly common in fisheries settings [31, 32] and has also been used to describe snow crab dynamics in the area of study [19]. The focus is usually upon biomass *B* and its normalized value, *b* = *BK*^−1^, rather than numerical abundance: *b*_*t*+1_ = *b*_*t*_ + Δ*b*_*t*_, where Δ*b*_*t*_ = *rb*_*t*_(1 − *b*_*t*_) − *Fishing*_*t*_*K*^−1^. Here, *Fishing*_*t*_ is scaled by *K* to make it non-dimensional and also in the interval (0,1); it is usually treated as an external factor (perturbation), measured without error. Of course, there is usually observation error and/or illegal removals that will get erroneously absorbed as a biological process; in this case by elevating the intrinsic rate of increase to compensate for the additional losses. This will have an effect of creating bias and uncertainty in related parameters (see below). Note also that the discretization to an annual basis is a significant assumption as the time span is large relative to the processes of most biota. This has the effect of averaging out sub-annual dynamics. This means temporal aliasing or discretization errors and censoring are introduced which ultimately increases process and observation errors (see subannual dynamics, below). From this *phenomenological* view, the minimal parameters required to estimate biological reference points, and the relative distance a system is from such reference points help delimit the status of a population [31, 32]. Values such as Maximum sustainable yield (*MSY* = *rK*/4) and the fishing mortality associated with such a yield (*FMSY* = *r*/2) are commonly used to help define some consistent landmarks of scale for use as management reference points to guide the implementation of a consistent *Precautionary Approach* for fishery exploitation [30].

Many approaches exist to estimate these model parameters. Currently, *Maximum Likelihood* approaches dominate due to their computational speed. However, it has been the author’s experience with this data that they do not navigate and optimize very high-dimensional parameter spaces reliably, especially in the delay differential equation models that we explore, below. This renders their utility in an operational setting, minimally useful. The related *Maximum A-Posteriori* solutions, where using the same optimization techniques but with the addition of “prior-like” constraints can result in marginally more stable results, but they still have tremendous difficulty with high dimensional parameter spaces and the associated numerous local optima/multiple equilibria.

Here we use Bayesian inference, as informative priors for these parameters are explicit and variance propagation of index variables can be accomplished by using them as priors to respective error terms. They facilitate parameter estimates that are stable and credible, given some prior knowledge of latent processes and observation models [32]. Previously, JAGS (Just Another Gibbs Sampler [33]) and STAN [34] were used to compute the posteriors via MCMC [19]. The latter uses the more efficient NUTS sampler which significantly speeds up the estimation process of these discrete difference equation models. Presently, we use **Julia** programming environment [20] and the supporting **Turing** library [21] for parameter inference and the **DifferentialEquations** library [22] for modeling delay differential equation models; they demonstrate efficiency and flexibility to simultaneously model and infer parameter values in such problems, relative to basic MCMC procedures due to use of automatic differentiation and heavily optimized solution engines.

Specifically, the latent (“real” but unobserved) biomass was assumed to have a *process model error* that is a recursive Gaussian or Normal distribution (*N*) with a standard deviation *σ*_*p*_ (Bolker 2008) such that:

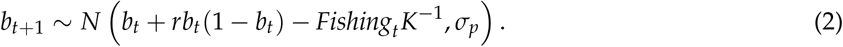

Here, “*∼*” indicates “distributed as”. Marginally informative priors were assumed: *r ∼ N* (1.0, 0.1), and *K ∼ N* (*κ*, 0.1 *· κ*). The prior mean of the carrying capacity (*κ* = [5.0, 60, 1.25], in kt, for the NENS, SENS, and CFA 4X, respectively; Figure 1) were based upon historical analyses with other analytical procedures, namely Geostatistical Kriging with External Drift and Generalized Additive Models [35, 12].

It is further assumed that the real unobservable (latent) process *b* is sampled with error (observation standard deviation *σ*_*o*_). The *observation error model* in most fishery applications is a simple multiplicative factor, often referred to as a “catchability” coefficient *q*. In this case we assume that: *q ∼ N* (1.0, 0.1), and so the *observation error model* becomes:

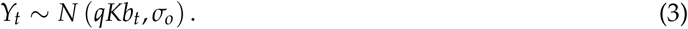

This means that the observed biomass indices *Y* are some constant fraction of the true abundance *b*_*t*_, across all sampling time (years, season) and locations. The recursive logistic process model (eq. 2) in tandem with the observation model (eq.3), we will refer to it as, “*Model 1*”.

From the graphical representation of the processes of *Model 1* (Figure 4), one can immediately see that there is no input term from outside of the system: it is a simple *phenomenological* self-loop model with two outputs (death and fishing). The term *rb* can be referred to as the (net) “birth process” [36, 37, 38], and represents a first order processes that increases *b* (via birth, growth, enhanced survival, improved food availability, movement, strong year class strength, etc.), through the single (static) parameter *r*. The mechanism is not identified and is implied to be some internal recycling of *b*. In reality, there are biological mechanisms involved and associated parameters that are not-static and non-stationary (in first and second order; across time and space). Growth in biomass or increase in numbers is seasonal and pulsed, due to the molting process and body mass increases, that progress at variable rates depending upon time of year and environmental variability. Across years, there are strong and weak year classes due to match-mismatch type processes [39, 40], and so also pulsed. These simple models cannot address these more realistic dynamics and so we obtain instead, an average state, with poor resolution of peaks and troughs and associated parameter estimates that are likely to be in error.

**Figure 4:**
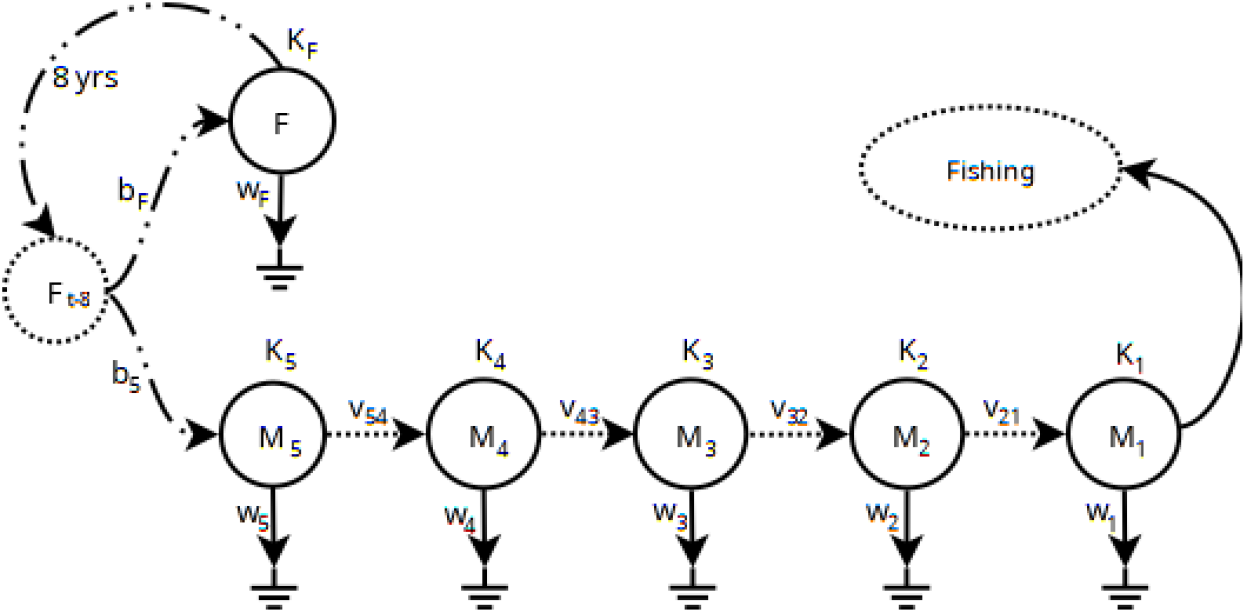
Graphical structure of the six-component Model 2 of snow crab with associated numerical flows. F indicates mature females, determined from size range and morphometric maturity. Associated with each component is a carrying capacity (K_i_), and a death rate w_i_, and molt rates v_ij_ ; and birth/survival rates b_i_. F_t−8_ is the state of mature females F from 8 years prior. Fishing is considered an external, deterministic sink imposed by external factors.

The strength of *Model 1* is simplicity; but this simplicity is also a weakness. As the biomass dynamics model is phenomenological; it is heuristic, without mechanism. For example, fishable biomass “creates” fishable biomass at a maximum exponential rate (r) to a limit of K. In reality, fishable biomass being mature male snow crab, does not produce more mature male snow crab; instead, it is the female mature population that creates the eggs and broods them to larvae, some fraction of which eventually mature into the fishable component. The dynamics and longevity of these females are very different from those of the males, often showing unbalanced sex ratios and differential utilization of habitat and of course complete unfished [9]. The rate of increase in biomass is not a simple exponential rate; instead, a certain number of fishable crab die from various causes (predation, disease, competitive fighting), a certain number of recruits terminally molt to a size in the fishable range to increase their numbers, they grow in weight as they eat, some fraction move in and out of a given area, and some are fished; these rates are not constant in time nor space and there are many lags in time. Fundamentally it is a numerical process. So *Model 1* from the perspective of “biomass dynamics” is less than satisfactory in terms of biological realism. However, in the heuristic perspective, it implies that the biomass in one year is related to the biomass in the previous year within some constraints, constraints that are only diffusely/indirectly related to the “real” mechanistic processes represented by individual-individual interactions. As such, they can be seen as a temporal autoregressive model.

These constraints unfortunately result in a model that is sometimes not responsive enough to large fluctuations in dynamics caused by intrinsic and/or extrinsic factors. For example, due to the pulse-like dynamics observed since the late 1990s, no recruits entered fishable components for a number of years, even though their abundance was low [12]. *Model 1* expected recruitment simply because biomass was low. This was an erroneous expectation due to the low numbers of pre-recruits that lasted for an extended period of time; the extreme warm bottom temperatures during this period and associated shifts in their spatial distributions; the increase in abundance of predators that followed which potentially reduced the strength of that recruitment. *Model 1* is too simple to express the expectations of such highly nonlinear pulse-like dynamics and interactions with environment and predators. In another example, there was an extreme warming event in 2021 that significantly altered and constricted the spatial distribution of snow crab [9]. Again a simple model such as *Model 1*, with static expectations of environmental conditions (*stationarity*) are not able to account for such effects.

In an attempt to *begin* addressing these issues, we entertain *Model 2* (Figure 4) which resolves some size groups, sex and maturity and permits time-varying viable habitat area (*non-stationarity*). *Model 2* is, therefore, an intermediately complex, marginally more structured and mechanistic (*ontological*) model relative to *Model 1*.

There is approximately an 8+ year period that is required for females *F*_*t*−8_ to produce the next generation of instar 9 females and males, denoted by *F*_*t*_ and *M*_5,*t*_, respectively. They represent crabs in the range of approximately 40 to 60 mm carapace width. The “birth” rates represent a combination of egg production and larval survival to instar 9+. If the population is stable, “birth” rates are expected to be similar in magnitude to overall death rates. Note that “stage” is defined based upon size-sex-morphometric traits and so misclassification is likely. The error in such knife-edge stage determinations can be substantial if growth and maturity schedules vary significantly. It is assumed, here, that in the aggregate, this has been stable in the survey record (1999 to 2022). This may of course be incorrect if persistent size/stage/age related mortality occurs due to exploitation/predation patterns and/or directional environmental change such as climate warming or food availability shifts.

Some fraction of instar 9 males transition to instar 10 (*M*_4_), and then 11 (*M*_3_) to 12 (*M*_2_) and 13+ (*M*_1_); each transition rate parameterized by *v*, with numeric indices identifying the instar pairs. Due to the knife edge-cut of stages, there will be misclassification issues which will be absorbed by these transition rates. For example, a small fraction of instar 11 crab (*M*_3_) will be large enough to be considered a fishable size (>95mm carapace width); but most will molt into instar 12 (*M*_2_); this would result in a reduced molt transition rate. *M*_2_ will represent a composite group of most crab that have entered fishable size but are still immature. Some fraction of this group will mature into the fishable component in the next year; the fractional nature will result in a smaller molt transition rate. Others will continue to molt to instar 13 and higher; for our purposes, all morphologically mature males of fishable size will be considered *M*_1_ and so it represents a range of different ages. This is especially the case as terminally molted crab can live for up to another 5 years; this aggregation will effectively decrease mortality rate of the group.

Survey sampling and estimation of instar 9 to 13+ (40 mm to 130+ mm carapace width) is reasonably informative. Earlier instars tend to have divergent habitat preferences relative to those of the fishable component and are known to be poorly/erratically sampled due to size approaching mesh size of sampling nets. This is because the snow crab surveys are primarily optimized for sampling the fishable component (> 95 mm carapace width), both in terms of mesh size and choices of sampling location. The utility of these additional compartments is that they also permit more stable parameter estimation and forward projections that are biologically more reasonable (mechanistic). The challenge is to keep track of much more information due to the increased realism and computational limits. Each state variable *U* = (*M*_1_, *M*_2_, *M*_3_, *M*_4_, *M*_5_, *F*), is scaled by their respective carrying capacity *K*., such that they are non-dimensional numbers: 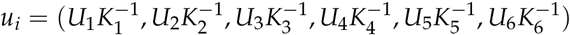. Thus, for example, in the molt transition process from *i* = 2 to *j* = 1, the instantaneous rate is:

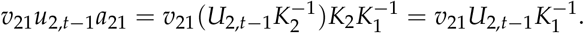

The multiplier 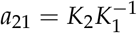 is required to convert normalized density of category 2 to that of category 1 via the ratio of the respective normalizing constants, *K*_*i*_. As with *Model 1*, first order birth/survival rates *b*_*i*_ are assumed: *β*_*i*_ = *b*_*i*_*u*_*i*_.

So far, most of the core model has been the same as *Model 1*, only applied to more components with some appropriate time lags. Where *Model 2* diverges from *Model 1* is by decomposing mortality into two components: the first component is a simple background mortality parameterized as a first order decay (*ω*_*i*_*u*_*i,t*_); the second component is mortality associated with fluctuations of viable habitat, parameterized as in the logistic equation as in *Model 1* as a second order process 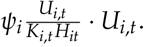. As such, it incorporates all density dependnent processes such as competition and interaction between stages. Note the congruence of this term with that of the 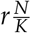 *· N* loss term of *Model 1* (see Fig. 4, eq. 1). With normalization to *u*, we obtain:

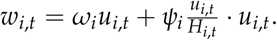

Here, *H* = *V*/max(*V*), represents the viable habitat surface area *V* normalized to a maximum value of 1, where the maximum is defined in the reference/focal time period. This modifies the carrying capacity proportionately and so *KH* can be seen as the “effective carrying capacity”, adjusting for viable habitat fluctuations. The assumption here is that when viable habitat area declines by some fraction *H*, so too does the effective carrying capacity, proportionately.

The inclusion of predator-prey and other ecological processes are therefore straightforward extensions as additional terms in this type of model formalism are well understood [41, 42, 43, 44]. Such extensions are planned for future research and evaluation.

The full set of delay differential equations are, therefore:

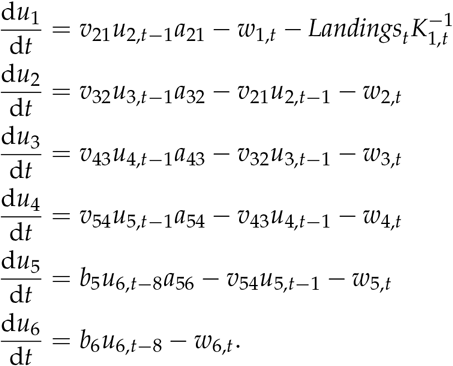

*Model 2* is, therefore, a more mechanistic (structured) extension of *Model 1* that brings to bear additional information available about the abundance of other size and sex classes and transition rates between them, and mortality rates that depend upon variations of numerical density in viable habitat surface area.

Fishery landings were converted to number based upon annual average weight of the fishable component estimated from surveys and discretized to a two week time step within each year. The latter was to avoid strong discontinuities which facilitates differential equation modeling and parameter inference. Survey-based indices of numerical abundance were estimated using a Hurdle process, where probability of observation and the numerical abundance of positive-valued observations were modeled as an extension of small-area analysis (Conditional AutoRegressive Models in space and time; see methods in [9, 26]. Modeled average weights, also analyzed using the same small-area space-time analyses were used to convert numbers back to biomass to make comparisons with *Model 1*.

In *Model 1*, the meaning of *q* is clear: a survey catches only some (fixed) fraction of the true abundance. This is a reasonable first approximation. However, what is implied is: first, this fraction is unchanging, applying equally in high abundance and low abundance years and areas, whether they are in the mating or molting season or not. Ignored are issues such as differential net saturation, aggregation behavior, even when bottom temperatures can force aggregation and dispersal away from or into other areas. The second issue implied by *q* is that when a survey fails to capture snow crab, that there truly is no snow crab in the area: an observation of zero is a true zero. ***This is clearly false***. Visual observation of trawl operation through video monitoring have shown that crabs are missed: when they are in a slight depression, when they are close to protruding rocks or on bedrock, when bottom topography is complex or simply by the gear jumping off of the bottom due to tidal currents. That is, sampling gear is biased. Furthermore, it is not just sampling gear that is problematic. Survey design is biased: in design, each areal unit is expected to be homogeneous in space AND time. External factors being factored out such as bottom temperatures, food availability and predation, aggregation behavior are not homogeneous, no matter how well designed a sampling design may be. Further, there is the notion of *trawlable bottom*: some areas cannot be sampled without tearing or losing nets. These areas are of course not sampled and only habitats that are easier to sample will be used and consequently, over represented. Though allocation may be *random* in design and theory, in application, it is at best, *almost random* and depending upon bottom type, usually biased.

As such, the observation model is also more complex in *Model 2*. The assumption of zero values in survey abundance indicating a true zero is a very strong assumption. This is especially tenuous when it is known that surveys sample many size ranges very poorly. The observation model of *Model 2* is, therefore, a simple linear regression. Of course, higher order terms and other covariates can and should enter into the observation model to account for potential survey and behavior induced bias. In this paper, this is mostly accomplished via the statistical abundance index model [9]. Some additional freedom is given in the observation model in that the prior for the intercept term *c*_*i*_ was informed by *µ*_*i*_, the fraction of the minimum observed value relative to the maximum value.

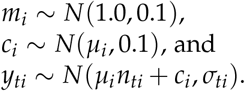

Here, *y*_*ti*_ = *Y*_*ti*_/max(*Y*_*i*_), constrains abundance to the interval (0, 1) and helps to confer better numerical properties (faster and more stable optimization, integration) in that variables are of similar magnitudes. Additionally, the variability of the observations was propagated into *Model 2* by assuming that the coefficient of variation of observations were modal estimates of the observation model using a Lognormal (LN) prior with a mode of log(CV) specific to each component *i* and survey time *t* and a SD of 0.25:

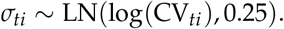

This works as *y*_*ti*_ are already normalized to (0,1). The other priors used for the *Model 2* are also informative based upon expected biological constraints and some very wide distributions for the variance components to provide well mixed posterior samples as determined by Effective Sample Size (given autocorrelation in samples within chains) and *Rhat* (convergence criterion across chains):

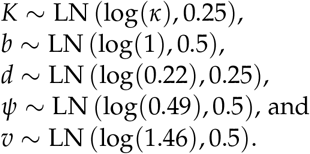

Fixed reference points analogous to the concepts of MSY and FMSY are not readily identifiable in *Model 2* as “production” is now dependent upon multiple system components each of which have externally imposed effective habitat viability trends and related natural and fishing mortalities (perturbations). That is, they are nonstationary, in the dynamical sense. It is, however, possible to compute a related concept, which we will call the *fisheries footprint*; it is computed as the difference in trajectories between the system state estimated for the fishable component with and without fishing. The biological parameters are estimated under conditions of fishing activity. In the absence of fishing, these parameters may be expected to be different, especially when fishing activity is extreme. In some cases, maturity and growth schedules can shift [45, 23, 46]. The assumption here is that extreme fishing activity has not occurred with sufficient pressure nor time to generate phenotypic or genotypic change. The difference in predicted trajectories between the fished and unfished conditions, therefore, provides a crude, first order estimate of the impact of fishing, without assumptions beyond that the model reasonably approximates reality. This *fisheries footprint* is placed on a relative scale from 0 to 1, by normalizing with the expected unfished abundance. As such, the *fisheries footprint* identifies the fraction of potential biomass that was reduced by fishery exploitation.

## 3 Results and Discussions

Estimates of abundance (biomass) of the exploited component are shown for each region for each model in Figure 5. In NENS, the discrete *Model 1* showed two troughs in (pre-fishery) abundance in 2005 and 2017. The surveyed index after adjustment for the observation model tends to be lower relative to the pre-fishery abundance, which is consistent with the removal of biomass during the fishing season. The exception to this pattern was in late 2013 and 2019 when strong recruitment was also observed. *Model 2* shows a similar pattern of fall fishable biomass, though with a reduced magnitude. The post 2019 period showed significant divergence relative to the survey index and especially in 2020 when no surveys were completed due to Covid-19 related disruptions. SENS also demonstrated a similar periodicity to NENS in both models (Figure 5). *Model 2* suggests a dampened time series, attributable to the dynamics of the recruitment being estimated to be flat (Figure 8). In CFA 4X, *Model 1* suggests an important decline following a peak in 2010. *Model 2* suggests a similar trajectory, though one that is much more variable. This area is subject to crab movement from the adjoining area as well as spatial aggregation due to extreme temperature conditions, elevated mortality from other predators and disease. *Model 2* also suggests overall abundance that is much lower than the very optimistic expectations of *Model 1*.

**Figure 5:**
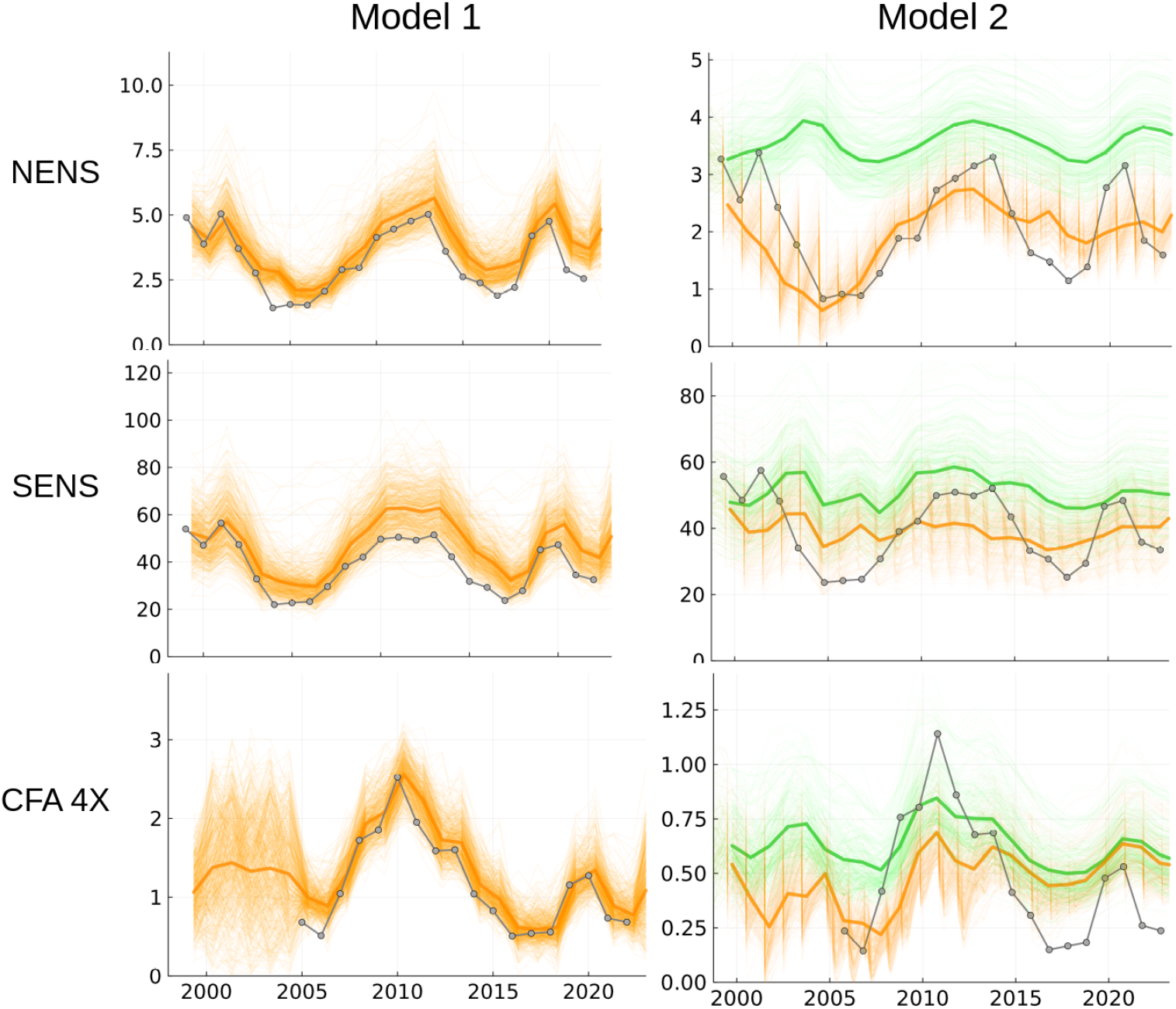
Posterior median biomass (dark orange) for each area of study and model. Posterior realizations of dynamics are shown to demonstrate variability of solutions. For Model 1 (left), pre-fishery posterior estimates of fishable biomass are shown. For Model 2, the posterior estimates of fishable biomass are shown in orange. Overlaid (gray) in both are survey abundance estimates (post-fishery, except prior to 2004) after correction for their respective observation models. Overlaid in green are posterior samples of abundance trajectories expected had there been no fisheries exploitation (green); and dark green identifies their respective means.

It is important to note that *Model 1* will project an increase in abundance, even if recruitment does not exist. It will only project a decline if abundance is above carrying capacity. This structural expectation is of course unrealistic. *Model 2*, which accounts for recruitment, navigates this problem a little more sensibly as an increase can only occur if recruitment exceed mortality. For example, it projects for CFA 4X, a decline, even in the absence of fishing. In SENS, abundance is projected to increase slightly in the absence of fishing. Only NENS was projected to have a rapid increase in abundance (Figure 5).

Another observation is that abundance may be overestimated by *Model 1*. For example, in CFA 4X where we have supporting information of very high natural mortality rates associated with predation and environmental variability, through the loss of strong year classes, abundance estimates are extremely optimistic. The same issues also exist for NENS, where strong adolescent crab year classes seem to disappear at a rate faster than might be expected, possibly due to predation. By extension, SENS also likely exhibits overly optimistic time trends, given the very strong year classes diminishing rapidly before entry into the fishable component. As a result of this potential overestimation of abundance, fishing mortality estimates by *Model 1* may in fact be overly low. *Model 2*’s solutions seem more reasonable given the supporting contextual information known of the different areas. The overall relative shapes of abundance and fishing mortality are, however, similar (Figures 5, 6).

**Figure 6:**
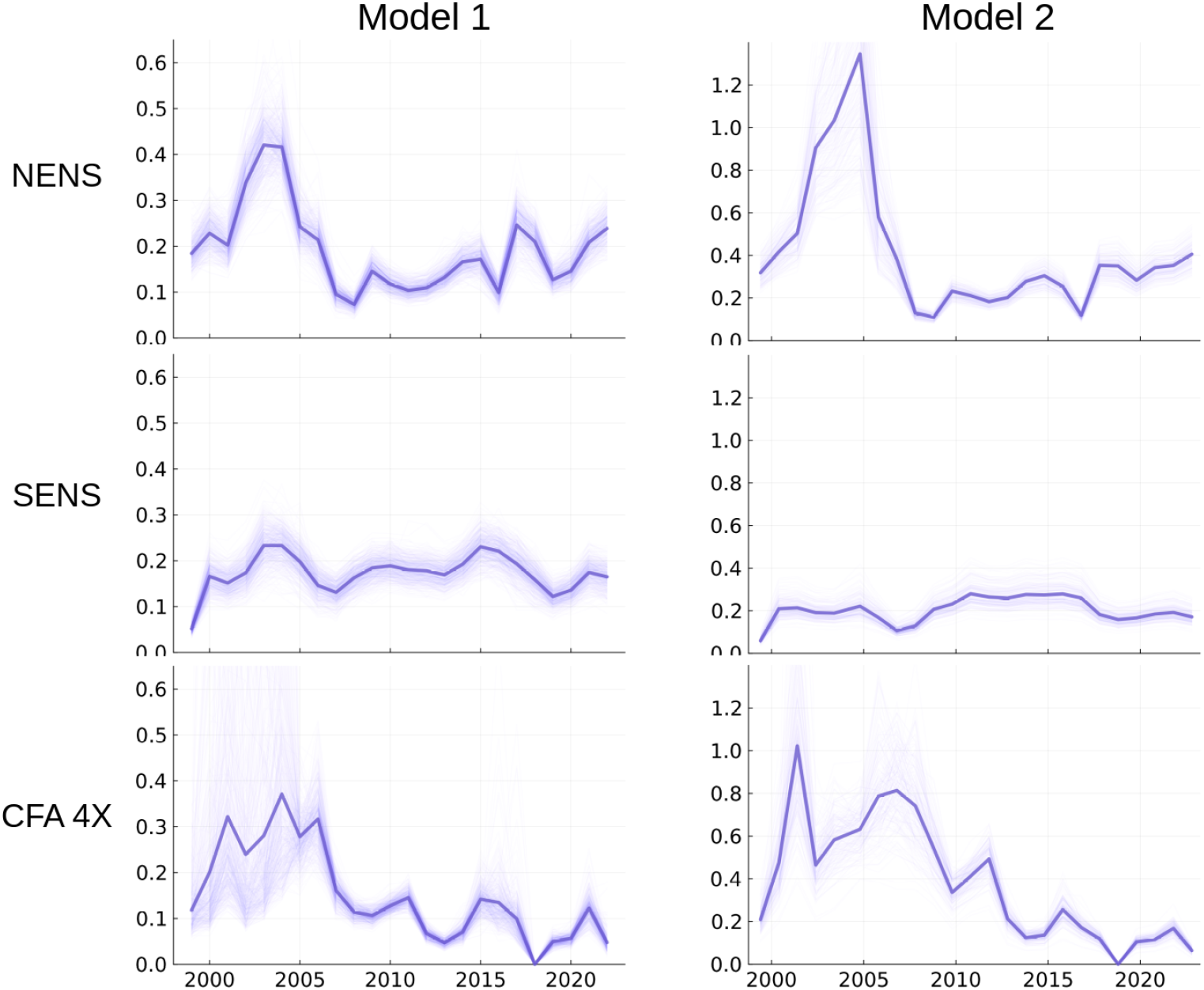
Instantaneous fishing mortality estimated from Model 1 and 2 across time. The overall patterns of fishing mortality are similar across models. However, the overall magnitudes are much higher in Model 2. Fishing in CFA 4X was closed in 2018 due to low abundance estimates and warm environmental conditions.

The continuous form of *Model 2* also permits us to infer the subannual dynamics of the snow crab. The saw tooth pattern of abundance (Figure 5) suggests that during the fishing season, the rate of exploitation far surpasses the slower rate of growth of the fishable component. It also identifies how important temporal aliasing might be if observations of landings or survey sample times are not properly accounted. The fishery footprint across the different areas (Figure 7) indicates that they are very similar to fishing mortality in overall form (as they should be, as the numerator, catch are the same). The former is on a scale that is the same as abundance and so simpler to understand than the exponential scale of an instantaneous rate. *Fishery footprint* also has a reasonably direct relationship to a question that is often asked in applied situations: *How much of an effect is fishing activity having?* In the case of the areas studied, the *fishery footprint* has declined to conservative and manageable levels (that is, a small fraction of the total available abundance) since the mid-2000’s. SENS, in particular, has been consistently sustainable in the historical record. In the presence of strong environmental variability, it will be necessary to continue to maintain a small *fisheries footprint* to reduce the susceptibility of the species such forces as has been the case for Atlantic cod and other collapsed fisheries [26]. This is especially the case as heavy exploitation in the long-term can result in reduced reproductive success of females; loss of dominance by invasion of habitat by competitors and even genetic/phenotypic selection for reduced size at maturity [47, 12].

**Figure 7:**
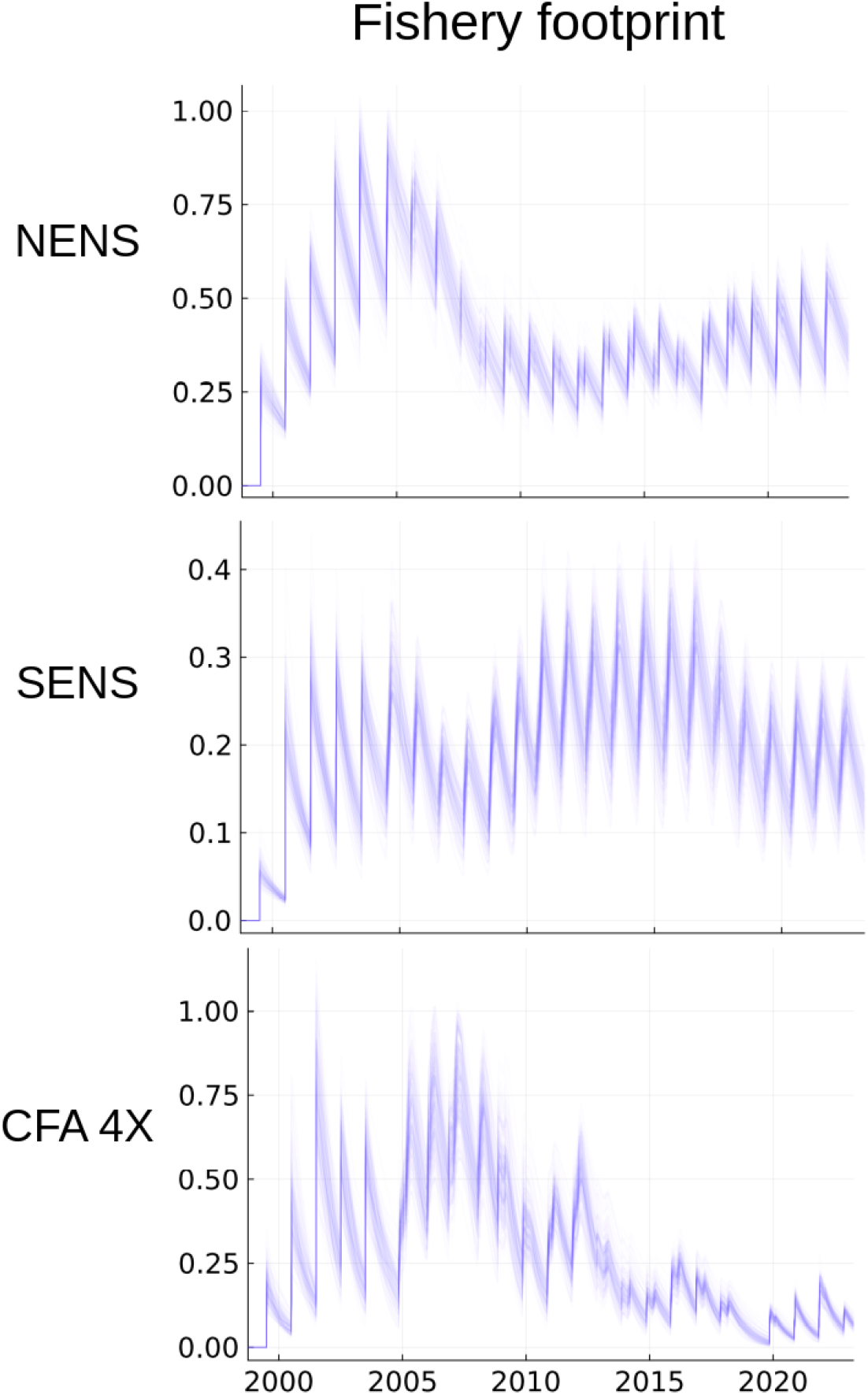
Posterior realizations of Fisheries footprint for each area of study derived from Model 2.

The first ever estimate of potential recruitment (*M*_2_) and mature female abundance (*F*) for the area in focus are shown in Figure (8). It shows that *Model 2* has difficulty tracking these components. However, peaks and troughs are identified and they seem to be coherent across areas. The lack of concordance with the observations suggest that other, as yet unmodeled processes likely need to be better accounted. The most likely missing processes include movement (inshore-offshore, shallow-deep) as well as predation, as the areas of study are not *well mixed* and show spatiotemporal structure and increases in the relative abundance of predators such as Atlantic Halibut [48]. Smaller areal units would also better parameterize the spatiotemporal heterogeneity observed in the latent biological processes known to occur in the area [12].

**Figure 8:**
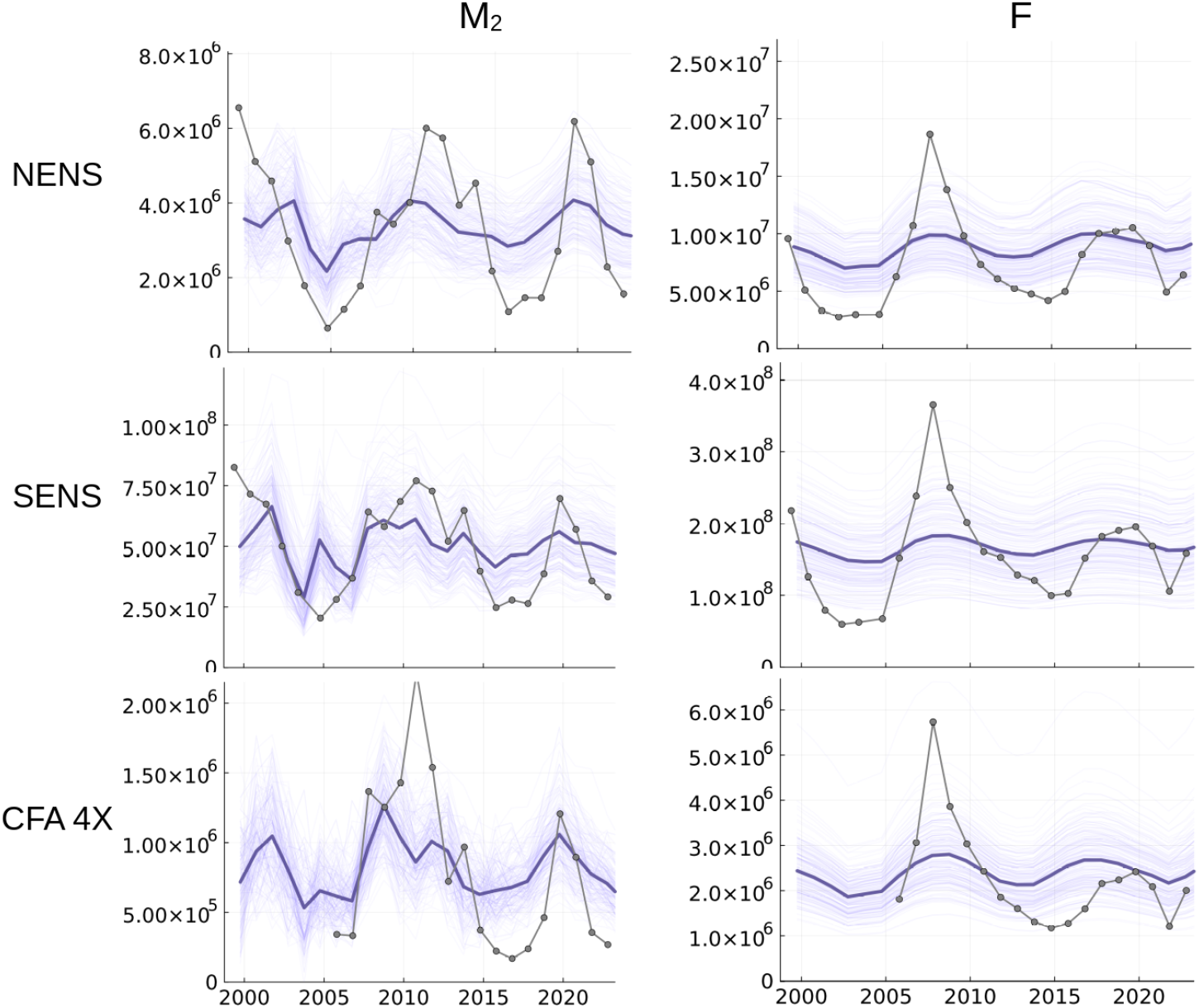
Model 2 posterior numerical abundance for M_2_ (recruits) and F (mature female). Overlaid (gray) are survey abundance estimates (post-fishery, except prior to 2004) after correction for observation models.

## 4 Conclusions

In studies of exploited marine animal populations, most models focus upon age structure [49]. Most also tend to be temporally discrete (annual) difference equation approximations. This is largely due to the annual cycle in temperate regions of exploitation, and data assimilation, survey and assessment cycles and historically due to limitations to computational capacity. With such discretizations, come various approximations and assumptions and ultimately potential error or bias from temporal (and spatial) aliasing. Space is treated as an externality or more usually, completely ignored, though of course, advection-diffusion differential equation models are readily formulated.

The continuous approach that we developed in this study addresses mechanisms in a structured manner. The additional model complexity succeeds in providing a similar and perhaps more reasonable understanding of snow crab population dynamics than the classical biomass dynamics model. It also permits a promising way forward towards describing the *fishery footprint* measured relative to potential, non-stationary abundance in the absence of fishing. It also identifies the importance of accounting for temporal dynamics with minimal aliasing in survey and landings, and the difficulties that may still need to be addressed: most notably the *non-well-mixed* nature of the areas studied show spatial and temporal structures that require further parameterization and development.

## References

[1] Yin-Juan Zhang, Bo Gao, and Xi-Wen Liu. “Topical and effective hemostatic medicines in the battlefield”. In: International Journal of Clinical and Experimental Medicine 8.1 (Jan. 2015), pp. 10–19. issn: 1940-5901.

[2] Babak Moghadas et al. “Novel chitosan-based nanobiohybrid membranes for wound dressing applications”. en. In: RSC Advances 6.10 (Jan. 2016), pp. 7701–7711. issn: 2046-2069. doi: 10.1039/C5RA23875G.

[3] J. C. Linden and R. J. Stoner. “Pre-harvest application of proprietary elicitor delays fruit senescence”. en. In: Advances in Plant Ethylene Research. Ed. by Angelo Ramina et al. Dordrecht: Springer Netherlands, 2007, pp. 301–302. isbn: 978-1-4020-6014-4. doi: 10.1007/978-1-4020-6014-4_65.

[4] Anna Tampieri et al. “Biologically inspired synthesis of bone-like composite: Self-assembled collagen fibers/hydroxyapatite nanocrystals”. en. In: Journal of Biomedical Materials Research Part A 67A.2 (2003), pp. 618–625. issn: 1552-4965. doi: 10.1002/jbm.a.10039.

[5] Aswani Poosapati et al. “Rechargeable Zinc-Electrolytic Manganese Dioxide (EMD) Battery with a Flexible Chitosan-Alkaline Electrolyte”. In: ACS Applied Energy Materials 4.4 (Apr. 2021), pp. 4248–4258. doi: 10.1021/acsaem.1c00675.

[6] T.P. Foyle, R.K. O’Dor, and R.W. Elner. “Energetically defining the thermal limits of the snow crab”. In: J. Exp. Biol. 145 (1989), pp. 371–393.

[7] B. Sainte-Marie and M. Lafrance. “Growth and survival of recently settled snow crab Chionoecetes opilio in relation to intra- and intercohort competition and cannibalism: a laboratory study”. In: Marine Ecology Progress Series 244 (2002), pp. 191–203.

[8] P.S. Kuhn and J.S. Choi. “Influence of temperature on embryo developmental cycles and mortality of female Chionoecetes opilio (snow crab) on the Scotian Shelf, Canada”. en. In: Fisheries Research 107 (Jan. 2011), pp. 245–252. issn: 0165-7836. doi: 10.1016/j.fishres.2010.11.006.

[9] J.S. Choi et al. “Temperature and depth dependence of the spatial distribution of snow crab”. en-CA. In: In prep (2022). doi: 10.1101/2022.12.20.520893. url: https://doi.org/10.1101/2022.12.20.520893.

[10] B. Sainte-Marie and F. Hazel. “Moulting and mating of snow crabs, Chionoecetes opilio, in shallow waters of the northwest Gulf of St. Lawrence”. In: Canadian Journal of Fisheries and Aquatic Sciences 49 (1992), pp. 1282–1293.

[11] M Comeau et al. “Growth, spatial distribution, and abundance of benthic stages of the snow crab, Chionoecetes opilio, in Bonne Bay, Newfoundland, Canada”. In: Canadian Journal of Fisheries and Aquatic Sciences 55 (1998), pp. 262–279.

[12] J.S. Choi. “Habitat Preferences of the Snow Crab, Chionoecetes opilio: Where Stock Assessment and Ecology Intersect”. In: Biology and Management of Exploited Crab Populations under Climate Change. Alaska Sea Grant, University of Alaska Fairbanks, 2011, pp. 361–376. isbn: 978-1-56612-154-5. doi: 10.4027/bmecpcc.2010.02. url: http://www.alaskaseagrant.org/bookstore/pubs/AK-SG-10-01.html.

[13] B Sainte-Marie. “Reproductive cycle and fecundity of primiparous and multiparous female snow crab, Chionoecetes opilio, in the Northwest Gulf of Saint Lawrence”. In: Canadian Journal of Fisheries and Aquatic Sciences 50 (1993), pp. 2147–2156.

[14] R. W. Elner and P.G. Beninger. “Multiple reproductive strategies in snow crab, Chionoecetes opilio: physiological pathways and behavioural plasticity”. In: Journal of Experimental Marine Biology and Ecology 193 (1995), pp. 93–112.

[15] Joel B. Webb et al. “Changes in Embryonic Development and Hatching in Chionoecetes opilio (Snow Crab) With Variation in Incubation Temperature”. In: The Biological Bulletin 213.1 (Aug. 2007), pp. 67–75. issn: 0006-3185. doi: 10.2307/25066619.

[16] M. S. M. Siddeek, E. Wade, and M. Moriyasu. “Proposed harvest control rule for managing the snow crab stock in the southwestern Gulf of St. Lawrence, Canada”. en. In: Fisheries Research 95.2 (Jan. 2009), pp. 268–279. issn: 0165-7836. doi: 10.1016/j.fishres.2008.09.032.

[17] Noel G. Cadigan, Elmer Wade, and Anders Nielsen. “A spatiotemporal model for snow crab (Chionoecetes opilio) stock size in the southern Gulf of St. Lawrence”. In: Canadian Journal of Fisheries and Aquatic Sciences 74.11 (Nov. 2017), pp. 1808–1820. issn: 0706-652X. doi: 10.1139/cjfas-2016-0260.

[18] J. Ianelli et al. GMACS. 2022. url: https://seacode.github.io/gmacs/.

[19] J.S. Choi and B.M. Zisserson. “Assessment of Scotian Shelf Snow Crab in 2010”. eng. In: Canadian Science Advisory Secretariat. Research Document 2011/100 (2011), pp. i–vii, 1–103.

[20] Jeff Bezanson et al. “Julia: A fresh approach to numerical computing”. In: SIAM review 59.1 (2017), pp. 65–98.

[21] Hong Ge, Kai Xu, and Zoubin Ghahramani. “Turing: a language for flexible probabilistic inference”. In: International conference on artificial intelligence and statistics, AISTATS 2018, 9-11 april 2018, playa blanca, lanzarote, canary islands, spain. 2018, pp. 1682–1690. url: http://proceedings.mlr.press/v84/ge18b.html.

[22] Christopher Rackauckas and Qing Nie. “Differentialequations.jl–a performant and feature-rich ecosystem for solving differential equations in julia”. In: Journal of Open Research Software 5.1 (2017).

[23] Jae S Choi et al. “Transition to an alternate state in a continental shelf ecosystem”. In: Canadian Journal of Fisheries and Aquatic Sciences 61.4 (Apr. 2004), pp. 505–510. issn: 0706-652X. doi: 10.1139/f04-079.

[24] Kenneth F. Drinkwater. “The response of Atlantic cod (Gadus morhua) to future climate change”. In: ICES Journal of Marine Science 62.7 (Jan. 2005), pp. 1327–1337. issn: 1054-3139. doi: 10.1016/j.icesjms.2005.05.015.

[25] J.S. Choi. “A framework for the assessment of snow crab (Chionoecetes opilio) in Maritimes Region (NAFO div 4VWX)”. In: DFO Can. Sci. Advis. Sec. Res. Doc. 2020/XXX (2020), viii + 68p.

[26] Jae S. Choi. “Reconstructing the decline of Atlantic Cod with the help of environmental variability in the Scotian Shelf of Canada”. en. In: In Prep (May 2022), p. 2022.05.05.490753. doi: 10.1101/2022.05.05.490753. url: https://www.biorxiv.org/content/10.1101/2022.05.05.490753v3.

[27] Håvard Rue et al. “Bayesian Computing with INLA: A Review”. en. In: arXiv:1604.00860 [stat] (Apr. 2016). arXiv: 1604.00860. url: http://arxiv.org/abs/1604.00860.

[28] Pierre François Verhulst. “Resherches mathematiques sur la loi d’accroissement de la population”. In: Nouveaux memoires de l’academie royale des sciences 18 (1845), pp. 1–41.

[29] Nicolas Bacaër. “Verhulst and the logistic equation (1838)”. en. In: A Short History of Mathematical Population Dynamics. London: Springer, 2011, pp. 35–39. isbn: 978-0-85729-115-8. doi: 10.1007/978-0-85729-115-8_6. url: https://doi.org/10.1007/978-0-85729-115-8_6.

[30] S. J. Smith et al. “Technical guidelines for the provision of scientific advice on the precautionary approach for Canadian fish stocks: Section 7–invertebrate species”. In: DFO Can. Sci. Advis. Sec. Res. Doc 117 (2012).

[31] M. B Schaefer. “Some Aspects of the Dynamics of Populations Important to the Management of the Commercial Marine Fisheries.” English. In: Bulletin of the Inter-American Tuna Commission 1 (1954).

[32] Renate Meyer and Russell B Millar. “BUGS in Bayesian stock assessments”. In: Canadian Journal of Fisheries and Aquatic Sciences 56.6 (June 1999), pp. 1078–1087. issn: 0706-652X. doi: 10.1139/f99-043.

[33] Martyn Plummer. JAGS: A program for analysis of Bayesian graphical models using Gibbs sampling. 2003.

[34] Stan Development Team. Stan Modeling Language Users Guide and Reference Manual, v 2.30.1. 2022. url: https://mc-stan.org.

[35] J.S. Choi, B.M. Zisserson, and A.R. Reeves. “An assessment of the 2004 snow crab populations resident on the Scotian Shelf (CFAs 20 to 24)”. In: DFO Can. Stock Assess. Res. Doc. 2005/028 (2005).

[36] Daniel T. Gillespie. “A rigorous derivation of the chemical master equation”. In: Physica A: Statistical Mechanics and its Applications 188.1 (Sept. 1992), pp. 404–425. issn: 0378-4371. doi: 10.1016/0378-4371(92)90283-V.

[37] Hong Qian and Lisa M. Bishop. “The Chemical Master Equation Approach to Nonequilibrium Steady-State of Open Biochemical Systems: Linear Single-Molecule Enzyme Kinetics and Nonlinear Biochemical Reaction Networks”. en. In: International Journal of Molecular Sciences 11.99 (Sept. 2010), pp. 3472–3500. issn: 1422-0067. doi: 10.3390/ijms11093472.

[38] Jaroslav Albert. “A Hybrid of the Chemical Master Equation and the Gillespie Algorithm for Efficient Stochastic Simulations of Sub-Networks”. en. In: PLOS ONE 11.3 (Mar. 2016), e0149909. issn: 1932-6203. doi: 10.1371/journal.pone.0149909.

[39] D. H. Cushing. “The Regularity of the Spawning Season of Some Fishes”. In: ICES Journal of Marine Science 33.1 (Nov. 1969), pp. 81–92. issn: 1054-3139. doi: 10.1093/icesjms/33.1.81.

[40] Joël M. Durant et al. “Climate and the match or mismatch between predator requirements and resource availability”. In: Climate Research 33.3 (2007), pp. 271–283. issn: 0936-577X.

[41] A. J. Lotka. “Analytical note on certain rhythmic relations in organic systems”. In: Proceedings of the National Academy of Science 6 (1920), pp. 410–415.

[42] L. von Bertalanffy. “The theory of open systems in physics and biology”. In: Science (New York, N.Y.) 111 (1950), pp. 23–29.

[43] E. R. Pianka. “On r- and K-selection”. In: The American Naturalist 104 (1970), pp. 592–597.

[44] P. J. Wangersky. “Lotka-volterra population models”. In: Annual Review of Ecology and Systematics 9 (1978), pp. 189–218.

[45] K.C.T. Zwanenburg et al. “Four Decadal changes in the Scotian Shelf Large Marine Ecosystem”. In: Large Marine Ecosystems. Vol. 10. Elsevier, Dec. 2002, pp. 105–150. isbn: 978-0-444-51011-2. doi: 10.1016/S1570-0461(02)80056-8.

[46] J. S. Choi et al. “Integrated assessment of a large marine ecosystem: a case study of the devolution of the eastern Scotian Shelf, Canada”. In: Oceanography and Marine Biology: An Annual Review 43 (2005), pp. 47–67.

[47] M. Comeau and G. Y. Conan. “Morphometry and gonad maturity of male snow crab, Chionoecetes opilio”. In: Canadian Journal of Fisheries and Aquatic Sciences 49 (1992), pp. 2460–2468.

[48] DFO. “Stock Status Update of Atlantic Halibut (Hippoglossus hippoglossus) on the Scotian Shelf and Southern Grand Banks in NAFO Divisions 3NOPs4VWX5Zc for 2020”. In: DFO Can.Sci. Advis. Sec. Sci. Resp. 2021/024 (2021).

[49] Terrance J. Quinn. “Ruminations on the Development and Future of Population Dynamics Models in Fisheries”. en. In: Natural Resource Modeling 16.4 (2003), pp. 341–392. issn: 1939-7445. doi: 10.1111/j.1939-7445.2003.tb00119.x.

